# The transcriptomic signature of physiological trade-offs caused by larval overcrowding in *Drosophila melanogaster*

**DOI:** 10.1101/2022.01.07.475433

**Authors:** Juliano Morimoto, Davina Derous, Marius Wenzel, Youn Henry, Hervé Colinet

## Abstract

Intraspecific competition at the larval stage is an important ecological factor affecting life-history, adaptation and evolutionary trajectory in holometabolous insects. However, the molecular pathways and physiological trade-offs underpinning these ecological processes are poorly characterised. We reared *Drosophila melanogaster* at three egg densities (5, 60 and 300 eggs/ml) and sequenced the transcriptomes of pooled third-instar larvae. We also examined emergence time, egg-to-adult viability, adult mass and adult sex-ratio at each density. Medium crowding had minor detrimental effects on adult phenotypes compared to low density and yielded 24 differentially expressed genes (DEGs) including several *chitinase enzymes*. In contrast, high crowding had substantial detrimental effects on adult phenotypes and yielded 2107 DEGs. Among these, upregulated gene sets were enriched in sugar, steroid and amino acid metabolism as well as DNA replication pathways, whereas downregulated gene sets were enriched in ABC transporters, Taurine, Toll/Imd signalling and P450 xenobiotics metabolism pathways. Overall, our findings show that larval overcrowding has a large consistent effect on several molecular pathways (i.e., core responses) with few pathways displaying density-specific regulation (i.e., idiosyncratic responses). This provides important insights into how holometabolous insects respond to intraspecific competition during development.

## Introduction

Intraspecific competition is an important evolutionary force leading to local adaptation, niche expansion and speciation through adaptive radiation [1,2]. Theoretical models have predicted – and empirical work has highlighted – that adaptation to novel resources is a key mechanism through which intraspecific competition affects niche expansion. For instance, theory predicts that high intraspecific competition in one niche or resource can lead to the invasion of underexploited niches by maladapted phenotypes [3] and that intraspecific competition modulates the degree of individual resource specialisation within populations [4]. Supporting evidence for these models comes from the work in *Drosophila melanogaster*, where populations experiencing high intraspecific competition levels adapt to cadmium-containing diets (a toxic compound) more rapidly than populations experiencing low intraspecific competition levels [5]. Similar patterns have been found in other species including tadpoles [6], sticklebacks [7], bees [8] and beetles [9] although in the latter species the pattern is less clear [10].

Holometabolous insects are particularly intriguing examples of these processes because intraspecific competition is likely to be the strongest at the larval stage rather than the adult stage. This is because of the underlying assumption is that insect larvae are often much less mobile than adults (see e.g., [11–13], but also [14]) and reliant primarily upon female oviposition site choices for feeding [15–17]. In these species, intraspecific competition at the larval stage triggers positive and negative responses that can carry-over to adult stage and modulate fitness. On the one hand, high intraspecific competition at the larval stage (‘overcrowding’) increases stress-tolerance through hormesis-like responses [18,19] which may result in longer adult lifespan [20]. Moreover, experimental evolution studies in *D. melanogaster* have also found that populations adapt to higher intraspecific competition at the larval stage by increasing thermal tolerance, feeding rates and tolerance to toxic compounds (e.g., urea) [21–24], which can be interpreted as a plausible proximate explanation for the diet niche expansion in high intraspecific competition populations [5]. Similar effects have been described in other drosophilids [25]. On the other hand, overcrowding constrains the availability of nutrients *per* capita, thereby constraining larval growth as well as adult morphology, behaviour and fitness [26,27]. In *D. melanogaster*, larvae developing in overcrowded conditions have unfavourable phenotypes that resemble those of individuals in protein-deficient larval diets [26], suggesting that overcrowding strengthens the competition for nutrients and limits the availability of protein which is essential for larval growth and adult reproduction. Moreover, overcrowding is also known to modulate the bacterial composition of the diet in which larvae are feeding which can affect pathogenicity of substrate, larval behaviour and life-history traits [28,29]. Similar effects were described in the tephritid fruit fly *B. tryoni* [30] and other insects (reviewed by [27]).

Although these examples highlight that physiological responses to overcrowding induce a range of phenotypic changes that can facilitate or hinder the survival and adaptation to local conditions, we know very little of the global gene expression patterns that underpin the responses to overcrowding. Previous target studies have examined molecular responses using only few candidate genes [18,19,22], which precludes our understanding of broad gene expression profiles that shape the entire suite of phenotypic responses to overcrowding. For example, Henry *et al*., [18] found higher relative expression of *Hsp70* and *Hsp40*, which is a sign of oxidative stress, in overcrowded conditions. Oxidative stress modulates a range of physiological responses that influence life-history traits, but we do not know the knock-on effects of higher oxidative stress on broader physiological processes. As a result, we lack a proper understanding of pleiotropic constraints and physiological trade-offs that can affect plastic and evolutionary responses to overcrowding and can in turn, modulate life-histories, niche expansion and ecological specialisation [31–33].

Here we address this knowledge gap by identifying for the first time the molecular basis of larval responses to overcrowding in a controlled experiment. We manipulated the intraspecific competition at the larval stage of *D. melanogaster* individuals using three larval density treatments [18,29] and investigated the global transcriptomic response to overcrowding. *Drosophila melanogaster* is the ideal model system to investigate the molecular responses to overcrowding for several reasons. First, *Drosophila* larvae are thought to be relatively immobile and, therefore, strongly influenced by interspecific competition [13,34]. Second, larval density modulates a range of fitness-related traits in both positive (e.g., hormesis-like) and negative (e.g., lower mating success) manners [18,19,22,29]. Third, responses to overcrowding modulate the strength of selection and can have intergenerational effects in *D. melanogaster* groups [35,36]. Lastly, responses to crowding are involved in increased larval competitive abilities that can result in niche expansion in evolutionary experiments in this model [5,20]. Table 1 shows our predictions for the highest density treatment based on the literature.

**Table 1.** Predictions.

| Predictions | Literature |
| --- | --- |
| Downregulation of metabolic pathways linked to dietary protein (e.g., TOR) due to protein limitation caused by high larval density (Prediction 1) | [26] |
| Upregulation of immunity due to density-dependent prophylaxis observed in other insects as well as upregulation of pathways related to survival (e.g., autophagy) *if* density-dependent responses increase survival (Prediction 2) | [20,37–39] |
| Upregulation of stress-related pathways (e.g., heat-shock proteins) as an hormesis-like response to overcrowding (Prediction 3) | [18,19] |
| No change or downregulation of pathways involved in reproduction (e.g., spermatogenesis) although some insects respond to high larval density by increasing testis sizes relative to body mass. | [27,40–42] |

## Material and Methods

### Larval density manipulation and adult phenotyping

We used an outbred laboratory population of *D. melanogaster* established from wild individuals collected in September 2015 in Rennes, Brittany (France). Fly stocks were maintained, and all experiments conducted, at 25°C and 70% relative humidity (12h light: 12h darkness) on standard food comprising inactive brewer’s yeast (MP Bio 029 033 1205, MP Bio, 80 g.L^−1^), sucrose (50 g.L^−1^), agar (Sigma-Aldrich A1296, 10 g.L^−1^) and supplemented with anti-mould Nipagin (Sigma-Aldrich H5501; 10% 8 mL.L^−1^). Adult flies from rearing stocks were allowed to lay eggs for less than 12 h on standard food supplemented with extra agar (15 g.L^−1^) and food dye. Eggs were collected with a paint brush, counted on moistened fabric and then transferred into new vials (50 mL; diameter = 23 mm) with 2.0 mL of standard food. We created three larval density treatments known to show contrasting phenotypes: L (low density; 5 eggs.mL^−1^), M (medium density; 60 eggs.mL^−1^) and H (high density; 300 eggs.mL^−1^) (see Henry et al. 2018, 2020). Egg manipulation was performed identically in all treatments and we standardised time spent under the stereomicroscope to 15 min to eliminate handling effects on the experiment. Eggs were allowed to hatch and larvae were allowed to develop until the wandering stage (third-instar larvae). At this precise point, we collected three replicates of 10 larvae per density treatment from independent vials (*N*_*larvae*_ *=* 90; *N*_*replicates*_ = 9) into cryo-tubes, which were snap-frozen in liquid nitrogen and stored at -80°C. Due to the lack of sex-specific morphological markers, larvae were selected without consideration of sex. We kept some vials unsampled (20, 2, and 2 respectively for conditions L, M and H) and allowed the flies to complete development until the adult stage in order to assess the viability, the development time, and the sex ratio. Recently emerged adults (<24h) were placed in single-sex vials with fresh food for three days before freezing at -20°C. Thirty males and thirty females per larval density treatment were then randomly collected, dried for one week at 60°C and weighed in a micro-balance (Mettler Toledo UMX2, Mettler Toledo, Greifensee, Switzerland; accurate to 1 μg). We examined effects of larval density on adult weight using generalised linear models (GLM) with Gaussian distribution followed by Student-Newman-Keuls post-hoc tests using the *agricolae* package [43]. Sexes were analysed separately and *P*-values were obtained from *F*-statistics.

### Larval transcriptome sequencing

Larval samples were ground to fine powder in 1.5 mL tubes placed in liquid nitrogen. Samples were mixed with 500 µL of lysis buffer (containing 1% β-mercaptoethanol) from Nucleospin® RNA extraction kits (Macherey-Nagel GmbH & Co. KG, Düren, Germany) and vortexed to complete homogenization. RNA extraction and purification were performed using Nucleospin® RNA columns (Macherey-Nagel) according to manufacturer’s instructions. Total RNA was eluted in 50 µL of RNase-free water. RNA was quantified and quality-checked using the NanoDrop ND-1000 spectrophotometer (NanoDrop Technologies, USA) and by running 200 ng of total RNA on 1% agarose gel. Quality control was also performed by Agilent Bioanalyzer Picochip (Agilent, Palo Alto, CA) which ensured high quality of all extractions (Figure S1). Illumina TruSeq stranded mRNA-Seq libraries were prepared and sequenced on an Illumina HiSeq 2500 instrument (Eurofins Genomics, Ebersberg, Germany), generating 100 bp paired-end reads.

The quality of the raw reads was examined in FASTQC 0.11 [44] and MULTIQC 1.7 [45]. Bases with a phred score below 20 and Illumina adapter read-through of at least 3 bp were trimmed from the 3’ end using TRIMGALORE 0.6.4 [46]. Trimmed reads were aligned to the *D. melanogaster* release 6 genome (GCF_000001215.4) using HISAT 2.1.0 [47] and alignments were processed using SAMTOOLS 1.9 [48]. To estimate the fraction of males in each larval pool, we obtained read coverage per chromosome using QUALIMAP 2.2.1 [49], standardised coverage to exome size as obtained from exon annotations using BEDTOOLS 2.28 [50] and calculated relative Y-chromosome coverage [Y/(X+Y)], expected to range from 0 (no males) to 0.5 (all males) [51].

### Differential gene expression

Read alignments were quantified against exon annotations and summarised at the gene level using FEATURECOUNTS 1.6.2 [52], considering only concordantly mapped read pairs and assigning multi-mappers as fractional counts to all mapping locations. Differential gene expression analysis among the three treatment groups was carried out with *DESeq2* 1.26.0 [53] in R 3.6.1 [54], shrinking fold-changes of low-count genes using the *apeglm* method [55]. To account for confounding differences in sex-specific trade-off responses [56] due to varying sex-ratios among pools, the model incorporated relative Y-chromosome coverage as a covariate. Differentially expressed genes (DEGs) were selected at an FDR-corrected [57]. *P*-value cut-off (*q*-value) of 0.05 and an absolute fold change cut-off of at least 1.5X. Overlaps of DEG lists between the three contrasts were inspected using intersection plots generated using *UpsetR* 1.4.0 [58]. Expression heatmaps were constructed using *gplots* in R [59]. To validate RNAseq data, expression level of 20 candidate DEGs was assessed with RT-qPCRs using the same (reverse-transcribed) RNA extracts (Table S1).

### KEGG pathway annotation and enrichment

We assigned KEGG pathway annotations to each gene using the KEGG pathway database (https://www.genome.jp/kegg/pathway.html) API as implemented in CLUSTERPROFILER 3.14.3 [60]. Gene names were converted to NCBI ENTREZ identifiers using the Bioconductor org.Dm.eg.db_3.10.0 database. We first examined the fold-change distributions of genes within each of the five main KEGG classes (Cellular Processes, Environmental Information Processing, Genetic Information Processing, Metabolism and Organismal Systems) and compared medians using the Kruskal-Wallis test. Second, we calculated the root-sum-square of fold changes within each KEGG class (regulation score). Third, we tested for significant functional overrepresentation of individual KEGG pathways against the *D. melanogaster* whole-genome background among sets of DEGs common across or unique to specific density contrasts. Significantly overrepresented KEGG pathways were selected at an FDR-corrected *P*-value cut-off (*q*-value) of 0.05. Finally, we constructed a network illustrating the overlap in genes between the significantly enriched KEGG pathways in Cytoscape version 3.7.1 with the plugin EnrichmentMap (default analysis type setting) [61,62].

## Results

### Larval density effects on adult phenotypes

Larval density affected the development dynamics, with individuals from the H treatment taking longer to reach adulthood than those from L and M (*t*-test, both adjusted *P* < 0.001) (Fig 1a). Viability from egg to adult was also impaired at higher densities, dropping from 91% [CI95: 86-94] in L to 84% [CI95: 78-88] in M and to 46% [CI95: 43-48] in H (Fig 1b). The sex-ratio of adult flies was slightly skewed towards females in all densities, ranging from 56% in L to 64% in H (Fig 1c). Larval density significantly affected both females’ (*F*_2,87_ = 261.54; *P* < 0.001) and males’ (*F*_2,87_ = 319.18; *P* < 0.001) dry body mass, whereby the H treatment generated adults that were significantly lighter than adults from L or M treatments (Fig 1d). These observations confirm that the H treatment was a resource-limiting and potentially stressful developmental environment for individuals, while the medium-density treatment had only minor effects.

**Figure 1:**
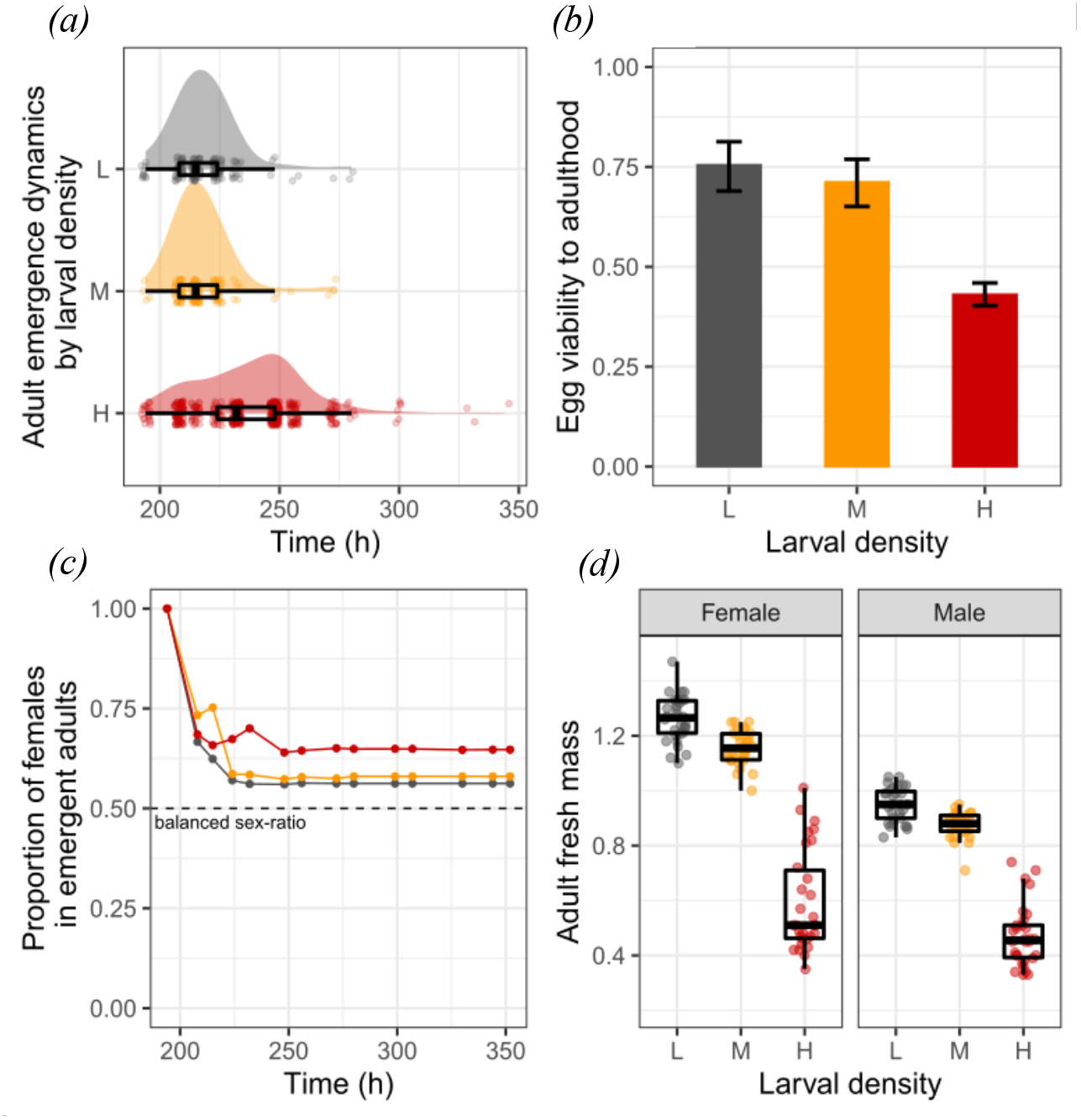
Effects of larval density on adult phenotypes. (a) adult emergence dynamics time. (b) egg to adult viability. (c) proportion of females in emergent adults. (d) body mass. Three larval densities were tested: L (low density; 5 eggs.mL^−1^; in grey), M (medium density; 60 eggs.mL^−1^; in orange) and H (high density; 300 eggs.mL^−1^; in red). Error bars in b) are 95% confidence intervals.

### Larval density effects on larval transcriptional patterns

We generated 18,988,311 to 28,154,203 high-quality read pairs per larval pool and quantified 82.3% to 86.9% of read alignments against gene annotations (Table S2). The reads covered 96.1% of the 17,759 annotated reference genes. Relative Y-chromosome coverage ranged from 0.09 to 0.54 (suggesting 2-10 males per pool; Table S2), and varied among density groups (*F*_2,6_ = 7.57; *P* = 0.02). Principal component analysis (PCA) on the variance-stabilised normalised read counts showed clear separation between the three larval density groups on the first axis (60% of variance explained; Fig 2a). The second PCA axis explained 26% of the variance and was significantly associated with standardised Y-coverage (*F*_1,7_ = 16.42; *P* = 0.005; Fig 2a). The lowest Y-coverage was observed at medium density, whereas samples at high density had highest Y-coverage but also large variation, overlapping with samples from low density (Fig 2a).

**Figure 2.**
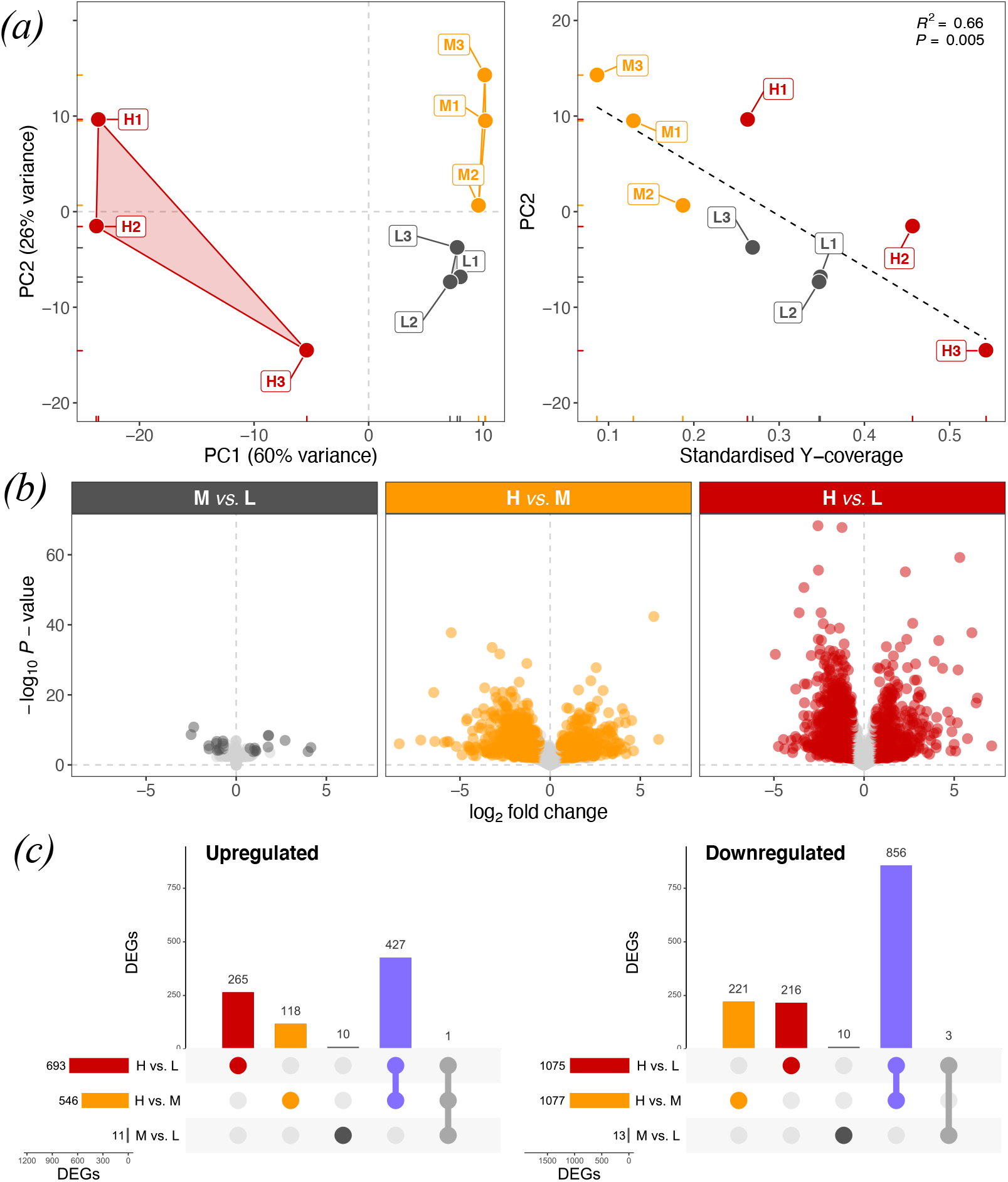
Effects of larval density on larval gene. (a) Left: Scatter plot of the first two principal components (PC1/PC2) of variance-stabilised normalised read counts among samples Right: Scatter plot and linear regression of PC2 against standardised Y-coverage (proxy for sex ratio in larval pool). The coefficient of determination (*R*^2^) and the *P*-value of the linear model are displayed on the top right. Samples are labelled and coloured by density treatment as in Figure 1. (b) Volcano plots summarising expression change (log_2_ fold change) and statistical significance (-log_10_ *P*-value) for each gene in medium/low (M *vs*. L), high/medium (H *vs*. M) and high/low (H *vs*. L) larval density contrasts. Significant DEGs (FDR <= 0.05, fold change >= 1.5X) are highlighted in dark grey (M *vs*. L), amber (H *vs*. M) or red (H *vs*. L). (c) intersection plots summarising overlap between upregulated (left) or downregulated (right) DEGs among the three contrasts. Four key intersections are highlighted in dark grey (M *vs*. L only), amber (H *vs*. M only), red (H *vs*. L only) and purple (DEGs common between H *vs*. M and H *vs*. L contrasts).

To account for this sex-specific effect, we incorporated standardised Y-coverage as a numeric covariate into the DGE model. The transcriptomic response was entirely consistent with phenotypic effects, whereby medium density had a weak effect and high density had a large effect on gene expression (Fig 2b). Fold changes within the three contrasts were tightly correlated with qPCR-based fold changes (*R*^2^ ranging from 0.95 to 0.97; Table S1). Only 24 differentially expressed genes (DEGs) were observed at medium density (M *vs*. L contrast), whereas 1623 and 1768 DEGs were observed at high density (H *vs*. M and H *vs*. L contrasts, respectively) (Fig S2, Table S3). The union of DEGs among these two high-density contrasts comprised 2107 DEGs, of which 1284 DEGs were shared (Fig S2). Considering overlaps among up- and down-regulated DEGs separately among all contrasts resulted in a “core” high-density crowding response of 427 up- and 856 down-regulated genes (Fig 2c, Table S3). For example, the *Hr38, ng1* and *ng3* genes were among the most strongly upregulated genes in the core crowding response (Table S3), and *ng2* was the only upregulated gene common across all three contrasts (Fig 2c). The H *vs*. M and H *vs*. L contrasts also contained 118 or 265 up- and 216 or 221 down-regulated unshared (“idiosyncratic”) DEGs, respectively (Fig 2c, Table S3). Although 19 of 24 DEGs at medium density were shared with other contrasts (Fig S2), the direction of regulation was opposite from that at high density for 20 of 24 DEGs (Fig 2c, Table S3). For example, *phu* was the most strongly upregulated gene at medium density but was among the most strongly downregulated genes at high density (Table S3).

### Biological pathway analysis of differentially expressed gene sets

We examined biological pathway annotations in each identified shared and unique gene set among density contrasts since these may represent distinct ecophysiological processes. The core crowding response (i.e., purple intersection in Fig 2c) comprised 1273 genes with strongly correlated fold changes between the H *vs*. M and H *vs*. L contrasts (*R*^2^=0.928; Fig 3a, Fig S2). Of these, 265 genes were annotated with KEGG pathways, most of which were from the “Metabolism” class. This class displayed by far the largest root-sum-square fold change among all five KEGG classes, for both up- and downregulated DEGs (Fig 3b). This strong regulation of metabolism was also observed for the idiosyncratic components of all three density contrasts (Fig 3c).

**Figure 3.**
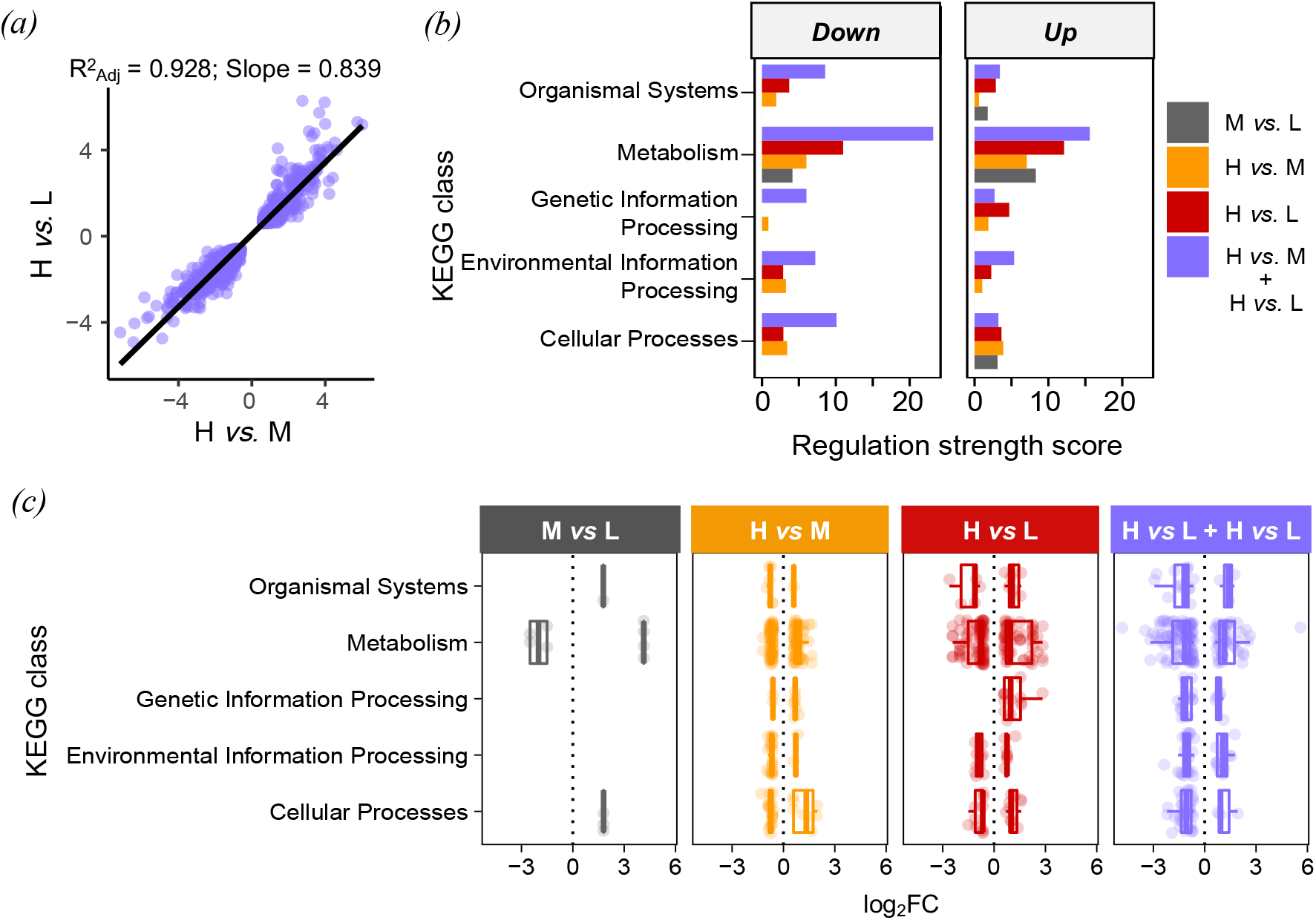
KEGG pathway regulation among DEGs responding to larval crowding. (a) correlation of log_2_ fold changes of DEGs in core crowding response between the two high-density contrasts (H *vs*. M and H *vs*. L). (b) Regulation strength scores (root-sum-square of fold changes) for each KEGG class based on the DEGs for the core crowding response and for unique DEGs in M *vs*. L, H *vs*. M, and H *vs*. L contrasts. (c) Genes log_2_FC organised according to KEGG class with boxplots superimposed for up- and down-regulated genes separately. Each dot represents a gene and dots are coloured according to contrast.

We next performed KEGG pathway enrichment (overrepresentation) analysis on the core crowding response and idiosyncratic responses. The upregulated DEGs in the core crowding response supported overrepresentation (FDR <= 0.05) of eleven pathways, including several metabolism pathways, *Purine metabolism* and *Steroid biosynthesis* (Fig 4, Table S4). The downregulated DEGs supported overrepresentation of *Sphingolipid metabolism, Lysosome, Taurine metabolism* and *Toll and Imd signaling* pathways (Fig 4, Table S4). The *Metabolic pathways* and *ABC transporters* pathways were significantly overrepresented both among up-and down-regulated DEGs (Fig 4, Table S4). There was little overlap in genes among these pathways, suggesting that these represent independent physiological responses (Fig 5). The idiosyncratic response to medium crowding (M *vs*. L) indicated downregulation of the *Amino sugar and nucleotide sugar metabolism* pathway, but no significant overrepresentation of any pathway among upregulated DEGs (Fig 4, Table S4). The two idiosyncratic responses to high crowding in the H *vs*. M and H *vs*. L contrasts included upregulation of *DNA replication* (both H *vs*. M and H *vs*. L) and *Mucin type O-glycan biosynthesis* (H *vs*. M. only), as well as downregulation of multiple *P450 xenobiotics metabolism pathways* (H *vs*. L only) (Fig 4, Table S4). Due to gene sharing with the P450 pathways, *Retinol metabolism, Ascorbate and aldarate metabolism* were also downregulated (Fig 6).

**Figure 4.**
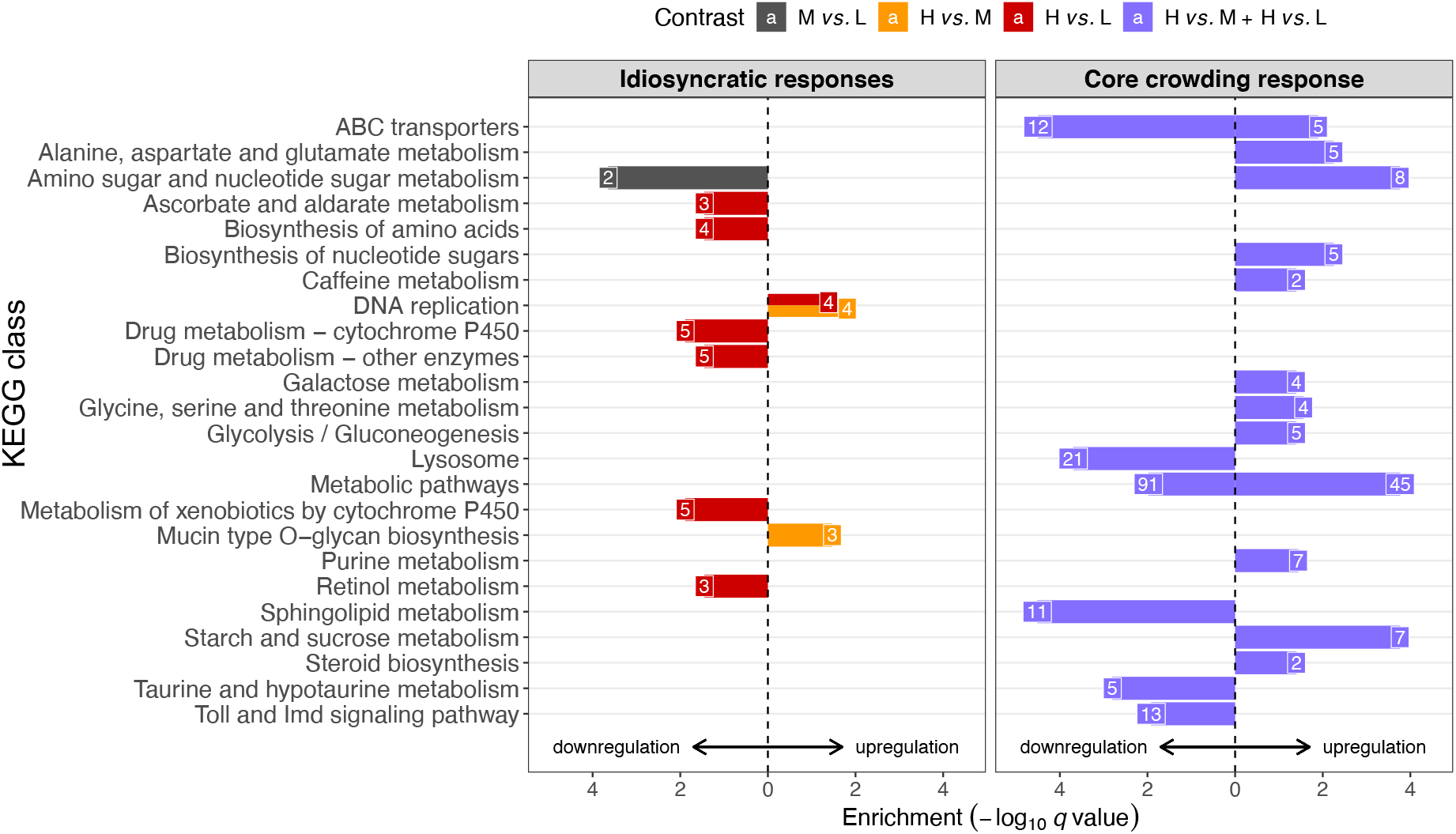
KEGG pathway enrichment in DEGs unique to individual density contrasts (idiosyncratic responses) or common in high/low and high/medium contrasts (core crowding response). The statistical significance (-log_10_ *q* value) is plotted for each KEGG pathway (*q* <= 0.05), whereby the direction of the bar indicates up- or downregulation of the underlying DEGs. The numbers of DEGs annotated with the enriched pathway are indicated at the end of each bar. Four key intersections are highlighted in dark grey (M *vs*. L only), amber (H *vs*. M only), red (H *vs*. L only) and purple (DEGs common between H *vs*. M and H *vs*. L contrasts).

**Figure 5.**
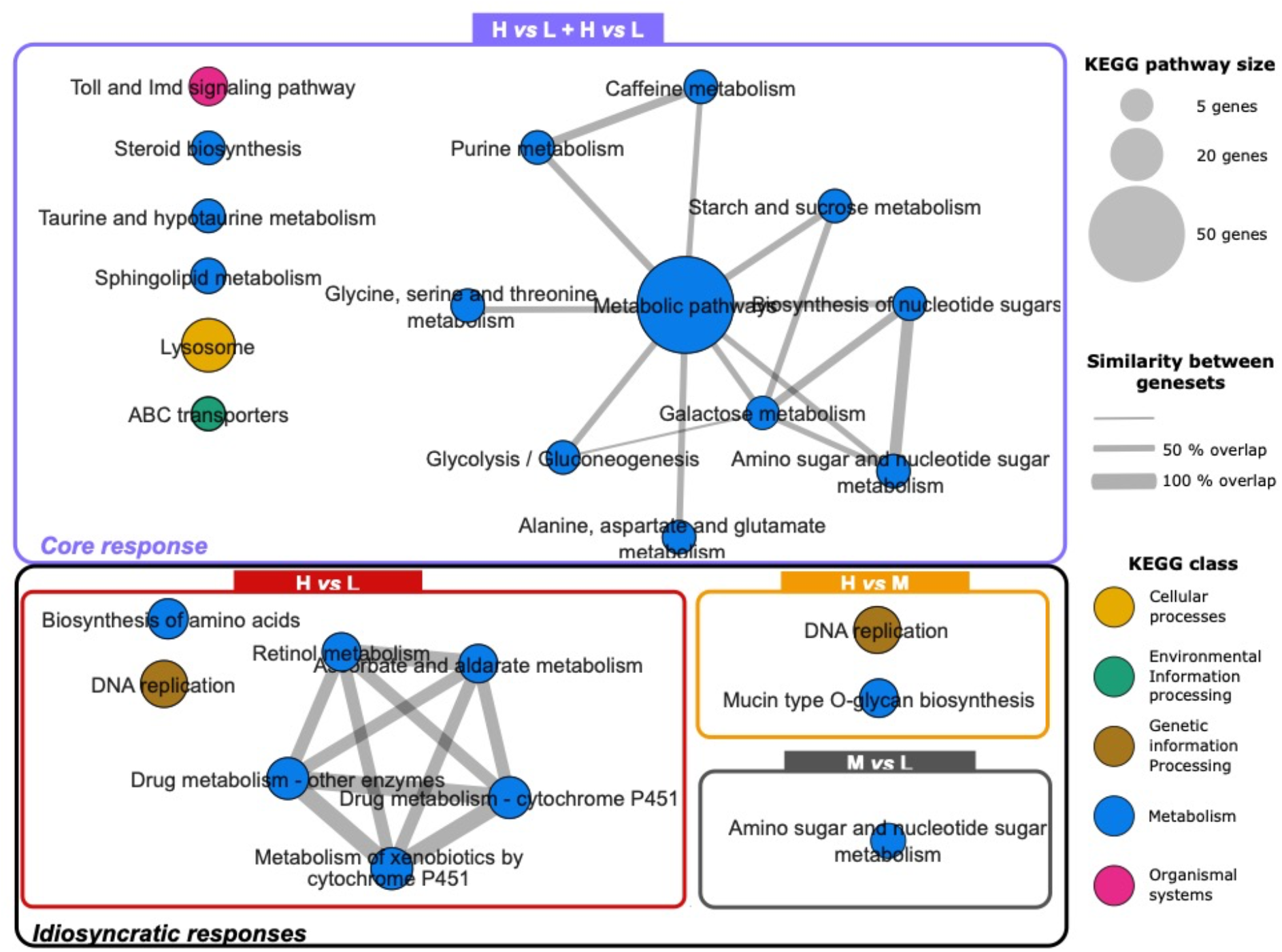
KEGG pathway similarity network for DEGs unique to three individual density contrasts (idiosyncratic responses) or common in H vs. M and H vs. L contrasts (core crowding response). The nodes represent the KEGG pathways significant at each comparison and the edges represent the similarity between the different KEGG pathways (i.e., overlap in genes). The size of the nodes represents the number of genes in the pathway with bigger nodes representing more genes. The width of the edges between the nodes represents the similarity coefficient. The thicker the edge the more genes they have in common. Pathways are coloured according to KEGG classification.

## Discussion

We present the first transcriptome-wide overview of physiological trade-offs caused by larval overcrowding in *Drosophila melanogaster*. Our results show that larval transcriptomic responses are consistent with the known negative adult phenotypic effects of larval overcrowding. While medium crowding had little effect on adult phenotypes and larval gene expression, high crowding had substantial phenotypic and transcriptomic effects, which were distinct from those at medium density. These data allow us for the first time to test predictions from previous targeted studies and provide novel candidate genes and physiological pathways for studying the ecological processes affected by larval overcrowding.

High crowding caused marked transcriptional changes in metabolic pathways and pathways related to growth and toxicity. This is consistent with decreased protein availability in the larval diet and an accumulation of toxic waste compounds, with knock-on effects for larval growth [18,26,29]. For instance, we found that steroid metabolism was upregulated, including lipases *Lip3* and *Lip4*. Interestingly Klepsatel *et al*., [26] reported that flies that had developed under high larval density had reduced protein content and were fatter than the flies from standard density. Likewise, Zwaan *et al*., [63] reported that the fat content of flies increased with increasing crowding. Hence it appears that crowding triggers a shift in metabolic pathways leading to triglyceride (fat) accumulation. This may be related to limiting dietary yeast/proteins in diet under crowded conditions [26,64]. This metabolic remodelling hypothesis was also suggested by metabolomic profiling data which revealed dissimilar metabotypes among larval densities which were mainly characterized by high concentrations of sugars and low amounts of amino acids at higher densities [18]. We also found that glycolysis, gluconeogenesis and galactose metabolism were upregulated probably as a by-product of elevated food uptake due to compensatory feeding to satisfy for protein requirements [26,64]. *Hr38* was among the most strongly upregulated genes, consistent with its role in regulating carbohydrate metabolism and glycogen storage of growing larval stage [65]. Moreover, purine metabolism was also upregulated as shown by the *uro* gene, which corroborates previous findings showing that *uro* is a key gene in the response to larval crowding [18]. Sphingolipid and taurine metabolism as well as lysosome and ABC transporters were downregulated suggesting that excretion and recycling pathways may be negatively impacted? in crowded conditions. Together, these results show that larval crowding modulates the response to low nutrition and high toxicity.

High crowding also caused strong downregulation of immunity via the Toll/Imd pathways involving two PGRPs (*PGRP-SD* and *PGRP-LA*) and *spätzle*, which resulted in downregulation of antimicrobial peptide expression including *DptA, Drs*, and *Drsl5*. This finding challenges the prediction that *Drosophila* larvae should upregulate immune function due to density-dependent prophylaxis as in other insects (e.g., [37,66]). However, it is possible that the larval densities used here, which are above those found in nature (see [34]), are above the larval densities which stimulates immunity. Nevertheless, the repression of transcripts involved in immunity found here is likely responsible for increased larval mortality in crowded conditions found here and elsewhere [27], which imposes a strong selective pressure for populations experiencing high larval densities. In *Drosophila*, populations adapted to larval overcrowding evolve pathogen-specific responses against gram-positive pathogens, suggesting that larval crowding might increase the risk of immune challenges by gram-positive pathogens in the population [38]. Crowding directly alters the nutritional properties of the diet, not only quantitatively but also qualitatively, with accumulation of metabolic wastes and decaying dead carcasses [18,29]. This in turn triggers marked changes in the bacterial composition of food with, for instance, the appearance of genus like *Pseudomonas* [29]. These bacteria are well known for their pathogenicity towards *Drosophila* [67]. Hence, the emergence of pathogens in food concurrently with a reduced immunity may be, at least in part, a driver of lethality at high larval density.

We also hypothesised that autophagy pathways would be upregulated in the core responses to crowding due to the reported increased lifespan in adults from crowded larval environments [20,26]. Our results do not directly support our predictions as our KEGG analysis did not detect an enrichment of the autophagy pathway. However, our analysis showed that the lysosome pathway, which is indirectly related to autophagy as well as directly involved in other biological processes, is downregulated in the core responses to high crowding. For instance, *LERP* is a receptor that is important lysosome vesicle trafficking as well as for eye development related to autophagy [68]. *LERP* knockdown results in lower body bass and increased starvation sensitivity in adult flies, which are traits associated with development in crowded environments [27]. Similarly, we found that *Npc2a* was downregulated, which is an important gene regulating the sterol metabolism and moulting in *Drosophila* larvae [69]. *Ppt2* was also downregulated and is a gene linked to neuronal development in *Drosophila* [70]. *Ppt2* knockdown showed abnormal neuronal development which could lead to impaired functions. This can help explain why *Drosophila* larvae developing in crowded conditions showed impaired learned visual recognition [71]. Autophagy is not only used for recycling damaged organelles or proteins; it also provides nutrients to maintain important cellular functions. Lipids including sphingolipids are increasingly recognized as key regulators of a series of critical cellular processes such as autophagy [72]. We found here that genes involved in sphingolipid metabolism were also downregulated in the core responses to larval crowding. Studies in flies have underscored the roles of sphingolipids in controlling lipid storage and response to nutrient availability [73]. For instance, changes in sphingolipid metabolic genes are part of the transcriptional response to sugar feeding of *Drosophila* larvae [74]. This study identified an acid sphingomyelinase (aSMase) and a ceramidase (CDase) both as being downregulated after sugar feeding, whereas fatty acid synthesis was upregulated. We also found that CDase was downregulated in response to crowding, as well as *schlank*, a ceramide synthase, and the ceramide kinase (*Cerk*) which catalyses phosphorylation of ceramide. Taken together, our results highlight sphingolipid metabolism is important during crowding, either for the regulation of cell death (autophagy) or for the metabolic regulation of nutrient scarcity.

Larval crowding was expected to trigger the expression of heat-shock proteins in a hormesis-like response [18,19]. However, our data did not show increased expression of heat-shock proteins at high density, suggesting that hormesis-like responses may not be as strong as previously considered. In a target gene expression study, Henry *et al*., [18] also failed to find strong upregulation of *hsp* family genes in M and H larvae, corroborating the present results. Thus, it appears that effects of crowding (beneficial or detrimental) are unrelated to heat-shock response. Nevertheless, high crowding downregulated the autophagy and lysosome pathways (i.e., recycling pathways) which could lead to a generalised stress response if damages in protein and other cellular components accumulate in individuals experiencing larval crowding. This accumulation of stress could underpin hormesis-like responses in adulthood. It is possible that our design, which sampled wandering larvae, selectively excluded larvae that accumulated higher stress levels as those were likely slower to pupate. This would explain the contradictory results presented here and the literature. Alternatively, it is possible that specific genes of the stress pathways are upregulated but not enough to be detected in our differential expression analysis. Future studies that directly manipulate larval stress levels will provide greater insights into the genes as well as overall transcriptional responses involved in organismal responses to stress, and will allow us to compare how those pathways are affected by larval crowding.

Larval crowding is also expected to affect reproductive processes, for example by increasing testis sizes relative to body mass [27,41,42]. Males and females developing in crowded larval environments are smaller and with lower mating and reproductive success, displaying higher investment in mating opportunities [27,75]. Moreover, phenotypic effects caused by larval crowding can persist through to next generations [35]. While our phenotypic data is consistent with these ideas, the transcriptomic data had no evidence of direct changes in reproductive pathways. One important limitation of our experimental design was that our larval pools had unknown sex-ratios and thus did not allow to specifically test for sex-specific effects on transcription. However, our sex-ratio estimates based on Y-chromosome coverage suggested that our pools differed in sex-ratio and that the transcriptomic response is affected by variation in sex-ratio. This is consistent with previous reports of sex-specific responses in reproduction to larval crowding [76–78] and warrants future efforts into ascertaining sex-specific crowding-associated trade-offs.

Three out of the four genes most strongly upregulated in response to high crowding were new glue genes *ng1, ng2, ng3*. These genes are normally transcribed from the beginning of the third instar and their expression is repressed later to allow the puffing stages transition to proceed in salivary gland chromosomes [79]. These intermoult puffs are regulated by ecdysone before the onset of metamorphosis, whereby an ecdysone peak can trigger key developmental events such as the cessation of feeding and the initiation of wandering of the larvae [80]. In *D. melanogaster* this ecdysone peak is also responsible for the repression of *ng1, ng2* and *ng3* [79,81]. In this study, we only sampled wandering larvae and the relative high expression of *ng* genes in crowded condition suggests a possible disturbance of ecdysone-mediated developmental processes (e.g., longer puffing stage transitions). This would be consistent with the observation that larval crowding affects the development dynamics, with individuals at high larval density taking longer to develop to adulthood than those from medium or low density.

The majority of the effects in molecular pathways belonged to the core responses to crowding, whereas the idiosyncratic responses to crowding revealed a relatively small number of pathways. For instance, medium crowding resulted in a small decrease in body mass and downregulation of *Cht4* and *Cht9* genes involved in the amino sugar and nucleotide sugar metabolism pathway. A further increase in larval crowding resulted in strong phenotypic effects as shown by a reduction in egg-to-adult viability and adult body weight (Fig 1), but upregulation of only repair enzymes of the DNA replication pathways (*Mcm* gens) and *Pgant* genes involved in mucin-type O-glycan biosynthesis pathways (H vs. M). Similar patterns were observed in the comparison between L and H, where the DNA replication pathway (*Mcm* and *DNApol* genes) were upregulated. Moreover, we also found that genes related to the metabolism of toxic compounds, including *Ugt* and *Gst*, were downregulated. In insects, *Ugt* genes are known to modulate the conjugation of glucose to eliminate toxic compounds and play a role in several physiological processes, including insecticide resistance [82,83]. Likewise, *Gst* are also involved in detoxification and insecticide resistance [84,85]. Therefore, H larvae have different susceptibility to toxic compounds relative to L larvae. Thus, increasing larval crowding has a consistent effect on molecular pathways above and beyond the magnitude of phenotypic effects (i.e., core response), and only specific pathways are regulated on a density-specific manner (i.e., idiosyncratic responses). Other Omics results also suggest that larval crowding generated physiological phenotypes that a are density-specific [18]. It is important to highlight that the *phurba tashi* (*phu*) gene, which is involved in hypoxia responses in *Drosophila* [86], showed a density-dependent expression pattern, whereby *phu* was strongly upregulated in M larvae but amongst the most downregulated in the H larvae. These density-specific physiological and molecular signatures may underlie hormetic responses triggered by crowding [18,19].

## Conclusion

We show that larval crowding induces changes in the transcriptomic profile of *Drosophila melanogaster* larvae. These responses are primarily related to metabolic pathways, but also include pathways related to growth and maintenance and toxicity. In particular, sugar, steroid and amino acid metabolism pathways as well as DNA replication, ABC transporters, Taurine metabolism, P450 xenobiotics metabolism and the Toll/Imd signaling pathways were identified as key pathways underpinning the molecular responses to larval overcrowding and thus, are candidates for future targeted studies. For instance, it will be important to characterise the specific role of the differentially expressed genes and pathways on the response to larval crowding across holometabolous insects, allowing for a comparative understanding of the molecular responses to intraspecific competition. Importantly, the molecular insights gained here help answer disparate questions in the field of developmental ecology, by shedding light into how molecular pathways respond to a stressful and competitive developmental environment. Overall, our study shows the molecular pathways underpinning larval responses to developmental environments and broadens our understanding of how holometabolous insects respond to intraspecific competition during development.

## Data accessibility

Raw sequence data and the processed gene-count matrix have been deposited at the NCBI Gene Expression Omnibus database [87] and are accessible through GEO Series accession number GSE193120

## Authors’ contributions

J.M: data curation, formal analysis, investigation, methodology, visualization, writing—original draft, writing - review and editing; D.D: data curation, formal analysis, methodology, resources, writing-review and editing; M.W: data curation, formal analysis, Investigation, methodology, validation, visualization, writing - review and editing; H.Y: Conceptualization, initial production of data, data curation, visualization, writing - review and editing; H.C: Conceptualization, methodology, initial production of data, formal analysis, supervision, funding, writing - review & editing. All authors gave final approval for publication and agreed to be held accountable for the work performed therein.

## Acknowledgements

We are grateful to EcoGeno Platform from UMS OSUR 3343 in Rennes for access to genomic facilities (2100 Bioanalyzer System and LC480 Roche qPCR System).

## Supplementary Materials

**Table S1**. Methods and Results of qPCR experiments to corroborate RNAseq data (separate excel).

**Table S2**. RNA-Seq QC and alignment statistics (separate excel).

**Table S3**. Full DGE results and intersections (separate excel).

**Table S4**. KEGG enrichment results (separate excel).

**Figure S1**. RNA electropherograms from Eukaryote Total RNA Pico assay

**Figure S2. Overlaps of differentially expressed genes (DEGs) in three density contrasts**. Patterns of gene expression in each intersection are visualised as heatmaps of log_2_ fold change (blue=downregulation; red=upregulation). M *vs*. L, H *vs*. M and H *vs*. L refers to medium-density versus low-density, high-density versus medium-density and high-density versus low-density respectively.

## References

1. Grant PR. 1972 Convergent and divergent character displacement. Biol. J. Linn. Soc. 4, 39–68.

2. Werner TK, Sherry TW. 1987 Behavioral feeding specialization in Pinaroloxias inornata, the “Darwin’s finch” of Cocos Island, Costa Rica. Proc. Natl. Acad. Sci. 84, 5506–5510.

3. Wilson DS, Turelli M. 1986 Stable underdominance and the evolutionary invasion of empty niches. Am. Nat. 127, 835–850.

4. Svanbäck R, Bolnick D. 2005 Intraspecific competition affects the strength of individual specialization: An optimal diet theory method. Evol. Ecol. Res. 7.

5. Bolnick DI. 2001 Intraspecific competition favours niche width expansion in Drosophila melanogaster. Nature 410, 463–466. (doi:10.1038/35068555)

6. Levis NA, Fuller CG, Pfennig DW. 2020 An experimental investigation of how intraspecific competition and phenotypic plasticity can promote the evolution of novel, complex phenotypes. Biol. J. Linn. Soc. 131, 76–87.

7. Svanbäck R, Bolnick DI. 2007 Intraspecific competition drives increased resource use diversity within a natural population. Proc. R. Soc. B Biol. Sci. 274, 839–844.

8. Fontaine C, Collin CL, Dajoz I. 2008 Generalist foraging of pollinators: diet expansion at high density. J. Ecol. 96, 1002–1010.

9. Agashe D, Bolnick DI. 2010 Intraspecific genetic variation and competition interact to influence niche expansion. Proc. R. Soc. B Biol. Sci. 277, 2915–2924.

10. Parent CE, Agashe D, Bolnick DI. 2014 Intraspecific competition reduces niche width in experimental populations. Ecol. Evol. 4, 3978–3990.

11. Jones RE. 1977 Search behaviour: a study of three caterpillar species. Behaviour 60, 237–259.

12. Hansen JD, Ludwig JA, Owens JC, Huddleston EW. 1984 Larval movement of the range caterpillar, Hemileuca oliviae (Lepidoptera: Saturniidae). Environ. Entomol. 13, 415–420.

13. Markow TA. 2015 The natural history of model organisms: the secret lives of Drosophila flies. Elife 4, e06793.

14. Goddard J, De Jong G, Meyer F. 2020 Unidirectional en masse larval dispersal of blow flies (Diptera: Calliphoridae). Food Webs 23, e00137.

15. Thompson JN. 1988 Evolutionary ecology of the relationship between oviposition preference and performance of offspring in phytophagous insects. Entomol. Exp. Appl. 47, 3–14. (doi:10.1111/j.1570-7458.1988.tb02275.x)

16. Larsson S, Ekbom B. 1995 Oviposition mistakes in herbivorous insects: confusion or a step towards a new host plant? Oikos, 155–160.

17. Gripenberg S, Mayhew PJ, Parnell M, Roslin T. 2010 A meta-analysis of preference– performance relationships in phytophagous insects. Ecol. Lett. 13, 383–393. (doi:https://doi.org/10.1111/j.1461-0248.2009.01433.x)

18. Henry Y, Renault D, Colinet H. 2018 Hormesis-like effect of mild larval crowding on thermotolerance in Drosophila flies. J. Exp. Biol. 221, jeb169342. (doi:10.1242/jeb.169342)

19. Lushchak O V et al. 2019 Larval crowding results in hormesis-like effects on longevity in Drosophila: timing of eclosion as a model. Biogerontology 20, 191–201. (doi:10.1007/s10522-018-9786-0)

20. Shenoi VN, Ali SZ, Prasad NG. 2016 Evolution of increased adult longevity in Drosophila melanogaster populations selected for adaptation to larval crowding. J. Evol. Biol. 29, 407–417.

21. Bubli OA, Imasheva AG, Loeschcke V. 1998 Selection for knockdown resistance to heat in Drosophila melanogaster at high and low larval densities. Evolution (N. Y). 52, 619–625.

22. Borash Teotónio, Rose, Mueller. 2000 Density-dependent natural selection in Drosophila: correlations between feeding rate, development time and viability. J. Evol. Biol. 13, 181–187. (doi:10.1046/j.1420-9101.2000.00167.x)

23. Shakarad M, Prasad NG, Gokhale K, Gadagkar V, Rajamani M, Joshi A. 2005 Faster development does not lead to correlated evolution of greater pre-adult competitive ability in Drosophila melanogaster. Biol. Lett. 1, 91–94. (doi:10.1098/2004.0261)

24. Mueller LD, Folk DG, Nguyen N, Nguyen P, Lam P, Rose MR, Bradley T. 2005 Evolution of larval foraging behaviour in Drosophila and its effects on growth and metabolic rates. Physiol. Entomol. 30, 262–269.

25. Nagarajan A, Natarajan SB, Jayaram M, Thammanna A, Chari S, Bose J, Jois S V, Joshi A. 2016 Adaptation to larval crowding in Drosophila ananassae and Drosophila nasuta nasuta: increased larval competitive ability without increased larval feeding rate. J. Genet. 95, 411–425. (doi:10.1007/s12041-016-0655-9)

26. Klepsatel P, Procházka E, Gáliková M. 2018 Crowding of Drosophila larvae affects lifespan and other life-history traits via reduced availability of dietary yeast. Exp. Gerontol. 110, 298–308. (doi:https://doi.org/10.1016/j.exger.2018.06.016)

27. Than AT, Ponton F, Morimoto J. 2020 Integrative developmental ecology: a review of density-dependent effects on life-history traits and host-microbe interactions in nonsocial holometabolous insects. Evol. Ecol. 34, 659–680. (doi:10.1007/s10682-020-10073-x)

28. Wong AC-N, Wang Q-P, Morimoto J, Senior AM, Lihoreau M, Neely GG, Simpson SJ, Ponton F. 2017 Gut Microbiota Modifies Olfactory-Guided Microbial Preferences and Foraging Decisions in Drosophila. Curr. Biol. 27. (doi:10.1016/j.cub.2017.07.022)

29. Henry Y, Tarapacki P, Colinet H. 2020 Larval density affects phenotype and surrounding bacterial community without altering gut microbiota in Drosophila melanogaster. FEMS Microbiol. Ecol. 96. (doi:10.1093/femsec/fiaa055)

30. Nguyen B, Ponton F, Than A, Taylor PW, Chapman T, Morimoto J. 2019 Interactions between ecological factors in the developmental environment modulate pupal and adult traits in a polyphagous fly. Ecol. Evol. 9. (doi:10.1002/ece3.5206)

31. Poisot T, Bever JD, Nemri A, Thrall PH, Hochberg ME. 2011 A conceptual framework for the evolution of ecological specialisation. Ecol. Lett. 14, 841–851.

32. Remold S. 2012 Understanding specialism when the jack of all trades can be the master of all. Proc. R. Soc. B Biol. Sci. 279, 4861–4869.

33. Kawecki TJ. 2008 Adaptation to Marginal Habitats. Annu. Rev. Ecol. Evol. Syst. 39, 321–342. (doi:10.1146/annurev.ecolsys.38.091206.095622)

34. Morimoto J, Pietras Z. 2020 Natural history of model organisms: The secret (group) life of Drosophila melanogaster larvae and why it matters to developmental ecology. Ecol. Evol. n/a. (doi:https://doi.org/10.1002/ece3.7003)

35. Morimoto J, Ponton F, Tychsen I, Cassar J, Wigby S. 2017 Interactions between the developmental and adult social environments mediate group dynamics and offspring traits in Drosophila melanogaster. Sci. Rep. 7. (doi:10.1038/s41598-017-03505-2)

36. Morimoto J, Pizzari T, Wigby S. 2016 Developmental environment effects on sexual selection in male and female Drosophila melanogaster. PLoS One 11. (doi:10.1371/journal.pone.0154468)

37. Siva-Jothy MT, Moret Y, Rolff J. 2005 Insect immunity: an evolutionary ecology perspective. Adv. In Insect Phys. 32, 1–48.

38. Kapila R, Kashyap M, Poddar S, Gangwal S, Prasad NGG. 2021 Evolution of pathogen-specific improved survivorship post-infection in populations of Drosophila melanogaster adapted to larval crowding. PLoS One 16, e0250055.

39. Schmid-Hempel P. 2003 Variation in immune defence as a question of evolutionary ecology. Proc. R. Soc. London. Ser. B Biol. Sci. 270, 357–366.

40. McGraw LA, Fiumera AC, Ramakrishnan M, Madhavarapu S, Clark AG, Wolfner MF. 2007 Larval rearing environment affects several post-copulatory traits in Drosophila melanogaster. Biol. Lett. 3, 607–610.

41. Johnson TL, Symonds MRE, Elgar MA. 2017 Anticipatory flexibility: larval population density in moths determines male investment in antennae, wings and testes. Proc. R. Soc. B Biol. Sci. 284, 20172087.

42. Morimoto J, Barcellos R, Schoborg TA, Nogueira LP, Colaço MV. 2021 3D imaging reveals potential anticipatory responses to increasing intraspecific competition in the <em>Drosophila</em> male reproductive system. bioRxiv, 2021.07.22.453343. (doi:10.1101/2021.07.22.453343)

43. De Mendiburu F. 2014 Agricolae: statistical procedures for agricultural research. R Packag. version 1.

44. Andrews S. 2010 FastQC: a quality control tool for high throughput sequence data. Version 0.11. 2. http://www.bioiniformatics.babraham.ac.uk/projects/fastqc.

45. Ewels P, Magnusson M, Lundin S, Käller M. 2016 MultiQC: summarize analysis results for multiple tools and samples in a single report. Bioinformatics 32, 3047–3048.

46. Krueger F. 2015 Trim galore. A wrapper tool around Cutadapt FastQC to consistently apply Qual. Adapt. trimming to FastQ files 516, 517.

47. Kim D, Paggi JM, Park C, Bennett C, Salzberg SL. 2019 Graph-based genome alignment and genotyping with HISAT2 and HISAT-genotype. Nat. Biotechnol. 37, 907–915. (doi:10.1038/s41587-019-0201-4)

48. Li H et al. 2009 The Sequence Alignment/Map format and SAMtools. Bioinformatics 25, 2078–2079. (doi:10.1093/bioinformatics/btp352)

49. Okonechnikov K, Conesa A, García-Alcalde F. 2016 Qualimap 2: advanced multi-sample quality control for high-throughput sequencing data. Bioinformatics 32, 292– 294. (doi:10.1093/bioinformatics/btv566)

50. Quinlan AR, Hall IM. 2010 BEDTools: a flexible suite of utilities for comparing genomic features. Bioinformatics 26, 841–842. (doi:10.1093/bioinformatics/btq033)

51. Palmer DH, Rogers TF, Dean R, Wright AE. 2019 How to identify sex chromosomes and their turnover. Mol. Ecol. 28, 4709–4724. (doi:https://doi.org/10.1111/mec.15245)

52. Liao Y, Smyth GK, Shi W. 2014 featureCounts: an efficient general purpose program for assigning sequence reads to genomic features. Bioinformatics 30, 923–930. (doi:10.1093/bioinformatics/btt656)

53. Love MI, Huber W, Anders S. 2014 Moderated estimation of fold change and dispersion for RNA-seq data with DESeq2. Genome Biol. 15, 550. (doi:10.1186/s13059-014-0550-8)

54. R Core Team. 2019 R: A language and environment for statistical computing. R Found. Stat. Comput.

55. Zhu A, Ibrahim JG, Love MI. 2019 Heavy-tailed prior distributions for sequence count data: removing the noise and preserving large differences. Bioinformatics 35, 2084– 2092. (doi:10.1093/bioinformatics/bty895)

56. Magwere T, Chapman T, Partridge L. 2004 Sex differences in the effect of dietary restriction on lifespan and mortality rates in female and male Drosophila melanogaster. Journals Gerontol. Ser. A 59, B3–B9. (doi:10.1093/gerona/59.1.B3)

57. Benjamini Y, Hochberg Y. 1995 Controlling the False Discovery Rate: A practical and powerful approach to multiple testing. J. R. Stat. Soc. Ser. B 57, 289–300. (doi:https://doi.org/10.1111/j.2517-6161.1995.tb02031.x)

58. Conway JR, Lex A, Gehlenborg N. 2017 UpSetR: an R package for the visualization of intersecting sets and their properties. Bioinformatics

59. Warnes MGR, Bolker B, Bonebakker L, Gentleman R, Huber W. 2016 Package ‘gplots.’ Var. R Program. tools plotting data

60. Yu G, Wang L-G, Han Y, He Q-Y. 2012 clusterProfiler: an R package for comparing biological themes among gene clusters. OMICS 16, 284–287. (doi:10.1089/omi.2011.0118)

61. Merico D, Isserlin R, Stueker O, Emili A, Bader GD. 2010 Enrichment map: a network-based method for gene-set enrichment visualization and interpretation. PLoS One 5, e13984.

62. Shannon P, Markiel A, Ozier O, Baliga NS, Wang JT, Ramage D, Amin N, Schwikowski B, Ideker T. 2003 Cytoscape: a software environment for integrated models of biomolecular interaction networks. Genome Res. 13, 2498–2504.

63. Zwaan BJ, Bijlsma R, Hoekstra RF. 1991 On the developmental theory of ageing. I. Starvation resistance and longevity in Drosophila melanogaster in relation to pre-adult breeding conditions. Heredity (Edinb). 66, 29–39. (doi:10.1038/hdy.1991.4)

64. Skorupa DA, Dervisefendic A, Zwiener J, Pletcher SD. 2008 Dietary composition specifies consumption, obesity, and lifespan in Drosophila melanogaster. Aging Cell 7, 478–490. (doi:10.1111/j.1474-9726.2008.00400.x)

65. Ruaud A-F, Lam G, Thummel CS. 2011 The Drosophila NR4A nuclear receptor DHR38 regulates carbohydrate metabolism and glycogen storage. Mol. Endocrinol. 25, 83–91.

66. Cotter SC, Hails RS, Cory JS, Wilson K. 2004 Density-dependent prophylaxis and condition-dependent immune function in Lepidopteran larvae: a multivariate approach. J. Anim. Ecol. 73, 283–293.

67. D’Argenio DA, Gallagher LA, Berg CA, Manoil C. 2001 Drosophila as a model host for Pseudomonas aeruginosa infection. J. Bacteriol. 183, 1466–1471.

68. Hasanagic M, Van Meel E, Luan S, Aurora R, Kornfeld S, Eissenberg JC. 2015 The lysosomal enzyme receptor protein (LERP) is not essential, but is implicated in lysosomal function in Drosophila melanogaster. Biol. Open 4, 1316–1325.

69. Huang X, Warren JT, Buchanan J, Gilbert LI, Scott MP. 2007 Drosophila Niemann-Pick type C-2 genes control sterol homeostasis and steroid biosynthesis: a model of human neurodegenerative disease. Development 134, 3733–3742.

70. O’Hern PJ. 2011 Investigation of Ppt2 RNAi Knock-down During Drosophila Neurogenesis. The FASEB Journal, 25, 753.5-753.5.

71. Slepian Z, Sundby K, Glier S, McDaniels J, Nystrom T, Mukherjee S, Acton ST, Condron B. 2015 Visual attraction in Drosophila larvae develops during a critical period and is modulated by crowding conditions. J. Comp. Physiol. A 201, 1019–1027. (doi:10.1007/s00359-015-1034-3)

72. Hebbar S et al. 2015 Ceramides And Stress Signalling Intersect With Autophagic Defects In Neurodegenerative Drosophila blue cheese (bchs) Mutants. Sci. Rep. 5, 15926. (doi:10.1038/srep15926)

73. Kraut R. 2011 Roles of sphingolipids in Drosophila development and disease. J. Neurochem. 116, 764–778. (doi:https://doi.org/10.1111/j.1471-4159.2010.07022.x)

74. Zinke I, Schütz CS, Katzenberger JD, Bauer M, Pankratz MJ. 2002 Nutrient control of gene expression in Drosophila: microarray analysis of starvation and sugar-dependent response. EMBO J. 21, 6162–6173. (doi:https://doi.org/10.1093/emboj/cdf600)

75. Wigby S, Perry JC, Kim Y, Sirot LK. 2016 Developmental environment mediates male seminal protein investment in D rosophila melanogaster. Funct. Ecol. 30, 410– 419.

76. Kapila R, Kashyap M, Gulati A, Narasimhan A, Poddar S, Mukhopadhaya A, Prasad NG. 2021 Evolution of sex-specific heat stress tolerance and larval Hsp70 expression in populations of Drosophila melanogaster adapted to larval crowding. J. Evol. Biol. 34, 1376–1385.

77. Vishalakshi C, Singh BN. 2008 Effect of environmental stress on fluctuating asymmetry in certain morphological traits in Drosophila ananassae: nutrition and larval crowding. Can. J. Zool. 86, 427–437.

78. Baldal EA, van der Linde K, van Alphen JJM, Brakefield PM, Zwaan BJ. 2005 The effects of larval density on adult life-history traits in three species of Drosophila. Mech. Ageing Dev. 126, 407–416. (doi:https://doi.org/10.1016/j.mad.2004.09.035)

79. D’Avino PP, Crispi S, Furia M. 1995 Hormonal regulation of the Drosophila melanogaster ng-gene. Eur. J. Entomol. 92, 259.

80. Dominick O, Truman JW. 1985 The physiology of wandering behaviour in Manduca sexta. II. The endocrine control of wandering behaviour. J. Exp. Biol. 117, 45–68.

81. Furia M, Digilio FA, Artiaco D, Giordano E, Polito LC. 1990 A new gene nested within the dunce genetic unit of Drosophila melanogaster. Nucleic Acids Res. 18, 5837–5841. (doi:10.1093/nar/18.19.5837)

82. Ahn S-J, Marygold SJ. 2021 The UDP-Glycosyltransferase Family in Drosophila melanogaster: Nomenclature Update, Gene Expression and Phylogenetic Analysis. Front. Physiol. 12, 300.

83. Lee S-W, Shono T, Tashiro S, Ohta K. 2005 Metabolism of PyracDofos in housefly, Musca domestica. J. Asia. Pac. Entomol. 8, 387–392.

84. Willoughby L, Chung H, Lumb C, Robin C, Batterham P, Daborn PJ. 2006 A comparison of Drosophila melanogaster detoxification gene induction responses for six insecticides, caffeine and phenobarbital. Insect Biochem. Mol. Biol. 36, 934–942. (doi:https://doi.org/10.1016/j.ibmb.2006.09.004)

85. Le Goff G, Hilliou F, Siegfried BD, Boundy S, Wajnberg E, Sofer L, Audant P, ffrench-Constant RH, Feyereisen R. 2006 Xenobiotic response in Drosophila melanogaster: Sex dependence of P450 and GST gene induction. Insect Biochem. Mol. Biol. 36, 674–682. (doi:https://doi.org/10.1016/j.ibmb.2006.05.009)

86. Mossman JA, Tross JG, Jourjine NA, Li N, Wu Z, Rand DM. 2017 Mitonuclear interactions mediate transcriptional responses to hypoxia in Drosophila. Mol. Biol. Evol. 34, 447–466.

87. Edgar R, Domrachev M, Lash AE. 2002 Gene Expression Omnibus: NCBI gene expression and hybridization array data repository. Nucleic Acids Res. 30, 207–210.

