## Supplementary Material for "The transcriptomic signature of physiological trade-offs caused by larval overcrowding in *Drosophila melanogaster*"

### Supplementary Tables

**Table S1.** Methods and Results of qPCR experiments to corroborate RNAseq data (separate excel).

**Table S2.** RNA-Seq QC and alignment statistics (separate excel).

**Table S3.** Full DGE results and intersections (separate excel).

**Table S4.** KEGG enrichment results (separate excel).

### Supplementary Figures

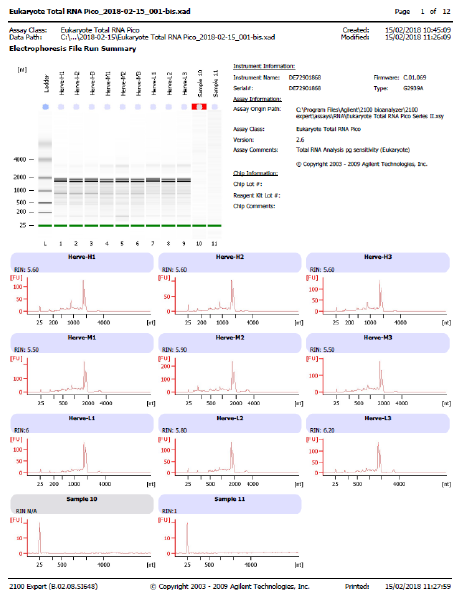

**Figure S1.** RNA electropherograms from Eukaryote Total RNA Pico assay

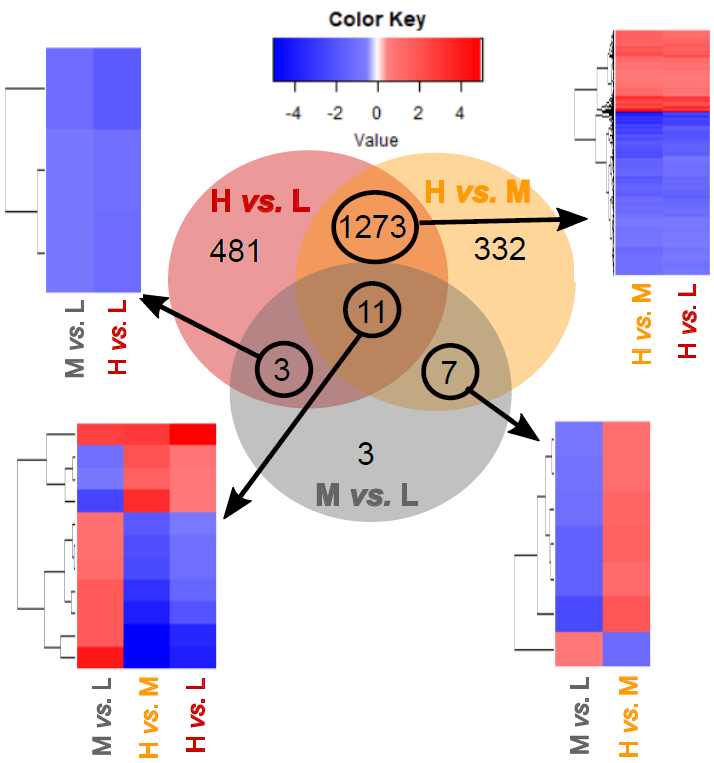

**Figure S2. Overlaps of differentially expressed genes (DEGs) in three density contrasts**. Patterns of gene expression in each intersection are visualised as heatmaps of log_2_ fold change (blue=downregulation; red=upregulation). M *vs.* L, H *vs.* M and H *vs.* L refers to medium-density versus low-density, high-density versus medium-density and high-density versus low-density respectively.
